# The microtubule nexus linking amyloid beta and tau: A simple and unifying theory for the underlying cause of Alzheimer’s Disease

**DOI:** 10.1101/2025.09.16.676664

**Authors:** Thomas A. Shoff, Maxence Derbez-Morin, Peishan Cai, Ryan R. Julian

## Abstract

Alzheimer’s disease (AD) is defined by cognitive decline in conjunction with accumulation of aggregated amyloid β (Aβ) and tau, yet existing models of AD fail to provide a simple connection between Aβ and tau. However, microtubules provide an intriguing nexus for pathological interactions between the two. Tau binds to microtubules and is critical to maintain their proper function. We demonstrate that Aβ also binds to microtubules with affinity comparable that of tau itself. We hypothesize that displacement of tau by Aβ leads to microtubule dysfunction and facilitates tau phosphorylation and aggregation. Importantly, in this model, aggregation of Aβ is not the primary cause of toxicity, which allows many of the apparent contradictions between Aβ pathology and cognition to be rationalized. This new model highlights the importance of both tau and Aβ and enables novel therapeutic and intervention strategies to be considered.

## Introduction

Alzheimer’s disease (AD) is the most common cause of dementia and has been the subject of many research endeavors worldwide. Despite this work, there are no effective cures or preventive measures against AD. Furthermore, the underlying cause of AD remains unclear and is the subject of considerable ongoing debate.^1-5^ Although frustrating, these observations are perhaps not surprising given that the disease is complex, as is the human brain and its workings, and the timescales for the occurrence and progression of the disease are measured in decades and years, respectively. To test and dismiss incorrect theories always requires time, but in the case of AD, researchers typically invest decades into a proposition before it can be properly evaluated. Pivoting toward new directions is therefore guarded by high barriers.

AD involves pathology that derives from two seemingly unrelated molecules, amyloid β (Aβ) and tau. These molecules form the aggregates in the brain that first allowed Alois Alzheimer to identify the disease. Not surprisingly, this propensity for aggregation has been the subject of significant interest. For example, Aβ aggregation (which occurs primarily in extracellular deposits) has been studied because many genetic variants associated with the processing of Aβ have been shown to cause early-onset AD in an autosomal-dominant fashion.^6^ Variants that increase the amount of Aβ or increase the aggregation propensity of Aβ accelerate the onset and progression of the disease. These observations drove the dominant theory about the cause of AD, the amyloid cascade hypothesis, which states that aggregation of Aβ is toxic and ultimately causes neuronal death. This basic theory has been presented in many variations and has evolved considerably over the years. However, the common theme is that aggregation of Aβ is the driving cause, and many therapies so focused have been attempted. The vast majority have failed, and the long-term efficacy of those currently remaining in use has yet to be determined.

Tau is also subject to aggregation in AD, with deposits appearing within neurons themselves.^7^ There are no known genetic mutations of tau that lead to AD, although mutations of tau are known and can lead to other tauopathies.^8^ Nevertheless, tau involvement is ubiquitous in AD. Critics of the amyloid cascade hypothesis have pointed out that tau pathology occurs earlier and correlates better with cognitive status than aggregation of Aβ.^7,9^ Indeed, the best predictors and indicators for AD are biomarkers derived from tau.^10-13^ The tau-centric hypothesis for the cause of AD focuses on aggregation of tau leading to neurotoxicity, with seeding of misfolded tau spreading the disease throughout the brain in a prion-like fashion.^14^

Both the Aβ- and tau-centric theories fail to account for the primary evidence used to support the opposing hypothesis, which appears to create the following irreconcilable contradictions:

1. If tau aggregation initiates AD pathology independently of Aβ, why do genetic mutations affecting Aβ strongly influence the disease?
2. If Aβ aggregation is the primary underlying cause, why does tau pathology occur first and serve as a better predictor of brain function? -and why have so many Aβ-driven therapies failed?

However, these observations can be reconciled if aggregation (of both Aβ and tau) is properly recognized as an effect of the underlying cause rather than the cause itself. Furthermore, the true cause should ideally involve direct interaction between tau and Aβ, thus immediately rationalizing the involvement of both molecules. Fortunately, a reasonable hypothesis that satisfies these criteria can be postulated, and it revolves around microtubules in neurons.

Microtubules are critical structures within neurons and are particularly important in axons where they enable transport of vital cellular components.^15^ Microtubules are inherently dynamic, with a variety of regulators that manage assembly/disassembly of alpha and beta tubulin into an open tube comprised of 13 proteins/ring.^16^ Tau is generally thought to stabilize microtubule assembly, particularly within neurons, and can also regulate transport by competitive binding relative to dynein or kinesin.^17^ Tau has also been reported to play a role in axonal elongation.^18^ Alternatively, it has been argued that tau may actually function to maintain microtubule lability (rather than stability).^19^ Regardless of the precise details, all evidence supports tau playing a crucial role in microtubule function within neurons.

Although the importance of tau for microtubule function is widely recognized, significantly less attention has been paid to interactions between Aβ and microtubules. Nevertheless, there are several experimental reports of binding between microtubules or tubulin proteins and Aβ. In the earliest work, agarose beads labeled with chimeric protein/Aβ1-42 constructs were used to pull down proteins from rat brain homogenates, revealing binding to tubulins.^20^ Pulldowns from human cell lines yielded the same result. In a similar study, Aβ 1-40 was immobilized on affigel that was used to make affinity columns. Soluble rat brain fractions run through these columns led to retention of half a dozen proteins including beta tubulin.^21^ A third extensive study examined many aspects of microtubule interactions with Aβ.^22^ Initial broad assays revealed binding between tubulin proteins and Aβ1-42, and subsequent targeted ELISA assays suggested a K_d_ of 400nM for the interaction. Other experiments showed that Aβ1-42 slowed the assembly and altered the ultimate morphology of microtubules. A fourth study used pelleting by centrifugation to coprecipitate proteins bound to Aβ1-42 fibrils, followed by identification with mass spectrometry.^23^ Forty total proteins were confidently identified as Aβ binders, including both alpha and beta tubulin. Aβ has also been reported to stabilize microtubules in the absence of tau, providing indirect evidence of a binding interaction.^24^ Finally, examination of the spatial distribution of Aβ within SK-N-SH cells revealed a high degree of colocalization with microtubules.^25^

In summary, a variety of past experiments suggest that various forms of Aβ1-40 and Aβ1-42 can bind to tubulins, which are the building blocks of microtubules. Although this interaction was noted in these previous works, the possibility that Aβ binding could impede the function of tau and therefore be crucial to the underlying cause of AD was not discussed.

Herein, we examine Aβ and tau microtubule binding domains for sequence homology, revealing high similarity. We further examine binding of fluorescently labeled Aβ1-40 and 1-42 to both individual tubulin proteins and microtubules. Fluorescence polarization (FP) confirms binding of monomeric Aβ to microtubules. Competition experiments monitored by fluorescence polarization with full length tau reveal binding comparable to Aβ. We propose that under conditions that lead to AD, binding of Aβ to microtubules interferes with tau binding, compromises microtubule functionality, and initiates tau toxicity. This framework for understanding AD allows many of the apparently contradictory observations related to tau and Aβ aggregation to be rationalized.

### Experimental

#### Materials

Porcine tubulin protein and taxol stabilized microtubules were purchased from Cytoskeleton. Amyloid beta 1-40 HiLyte Fluor 488-labeled and Amyloid beta 1-42 HiLyte Fluor 488-labeled were purchased from Anaspec. Wide bore pipette tips were obtained from Avantor Sciences. Paclitaxel (Taxol), EGTA, and anhydrous DMSO were from Sigma Aldrich. Recombinant human 2N4R Tau (rhTau) was purchased from rPeptide. PIPES, MgCl_2_, NaOH, NH_4_OH, and Optima grade water were obtained from Thermo Fisher Scientific. 384-well microplates (Polystyrene, black, F-bottom) were purchased from Greiner Bio-One.

#### Buffer preparation

For all experiments using monomeric tubulin, tubulin buffer was prepared with final concentrations of 80 mM PIPES, 2 mM MgCl_2_, and 0.5 mM EGTA in H_2_O. For all experiments using microtubules, microtubule buffer was prepared with final concentrations of 15 mM PIPES and 1 mM MgCl_2_ in H_2_O. Taxol was reconstituted using anhydrous DMSO to a concentration of 2 mM. For every 10 mL of microtubule buffer, 100 µL of taxol was added immediately prior to use.

#### Aβ disentanglement

Aβ peptides were disentangled as previously reported.^26^ Briefly, 0.1 mg Aβ-HiLyte samples were separately reconstituted in 200 µL of 10% NH_4_OH and incubated for 10 minutes at room temperature. Samples were then sonicated for 5 minutes and lyophilized to dryness. To prevent aggregation, Aβ samples were next reconstituted in 60 mM NaOH, aliquoted, and stored at -80°C. Prior to use, samples were thawed and reconstituted in the appropriate buffer to 50 nM. Aβ concentration was confirmed via UV-Vis spectroscopy using absorbance at 503 nm and an ε of 71,490 M^-1^cm^-1^.

#### Fluorescence polarization

All tubulin and microtubule samples were reconstituted in their appropriate buffers and serially diluted. Any solutions containing microtubules were handled using wide-bore pipette tips to ensure microtubule integrity and prevent clogging. Aβ-HiLyte solution was added to a final concentration of 10 nM to yield sufficient fluorescence intensity while remaining monomeric. Samples were loaded into 384-well plates, lightly tapped to remove any bubbles, incubated for 15 minutes, and loaded into the BioTek Synergy H1 plate reader. For all selected wells, triplicate FP readings were collected using an excitation filter of 485 ± 20 nm and an emission filter of 535 ± 20 nm. Any experiments using microtubules were collected using a time course mode where triplicate FP readings were collected every 2 hours for 24 hours. Polarization was calculated as:

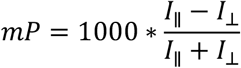

where *I*_∥_ corresponds to parallel intensity and *I*_⊥_ corresponds to perpendicular intensity. All polarization values were normalized to a blank containing only Aβ-HiLyte prior to fitting. K_D_ values were obtained by fitting the change of polarization as a function of tubulin or microtubule concentration to a quadratic equation:

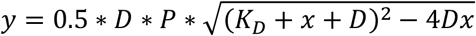

where y is fluorescence polarization, D is the concentration of Aβ-HiLyte, x = the concentration of tubulin or microtubules, and P is the maximal polarization.

For competitive binding experiments, Aβ and microtubules were incubated for 24 hours at room temperature to allow for binding. rhTau was reconstituted in water, diluted to an appropriate concentration in microtubule buffer, and spiked in at a final concentration of 2 µM. FP was collected as above. For co-equilibration experiments, Aβ, microtubules, and rhTau were incubated together and FP was monitored. To examine the relative change in polarization due to addition of tau and account for slight differences in microtubule concentration, ΔmP was normalized to 1 for the initial time points for all samples.

## Results

### Tau binding domain versus Aβ homology

Considerable existing data indicates that Aβ can bind to microtubules, suggesting that it may share sequence similarity with tau. The four recognized and two putative binding domains of tau are similar in size to Aβ (∼30 residues versus ∼40 residues). The potential similarity between Aβ and these binding domains was evaluated with several sequence homology models, as illustrated in Fig. 1. Cobalt suggests high sequence homology for 22 residues (Fig. 1a).^27^ Furthermore, the similarity between the various tau binding domains is nearly identical to that with Aβ. Similar results were obtained with M-Coffee (Fig. 1b), particularly for comparison between Aβ, P2 and R’.^28^ Espript 3.0 is a more structurally oriented homology search, which also finds similar homology levels between all of the tau binding domains and Aβ (Fig. 1c).^29^ It is also known that both Aβ and tau microtubule binding domains form amyloid fibrils, further suggesting that shared structural propensities exist for these sequences.

**Figure 1.**
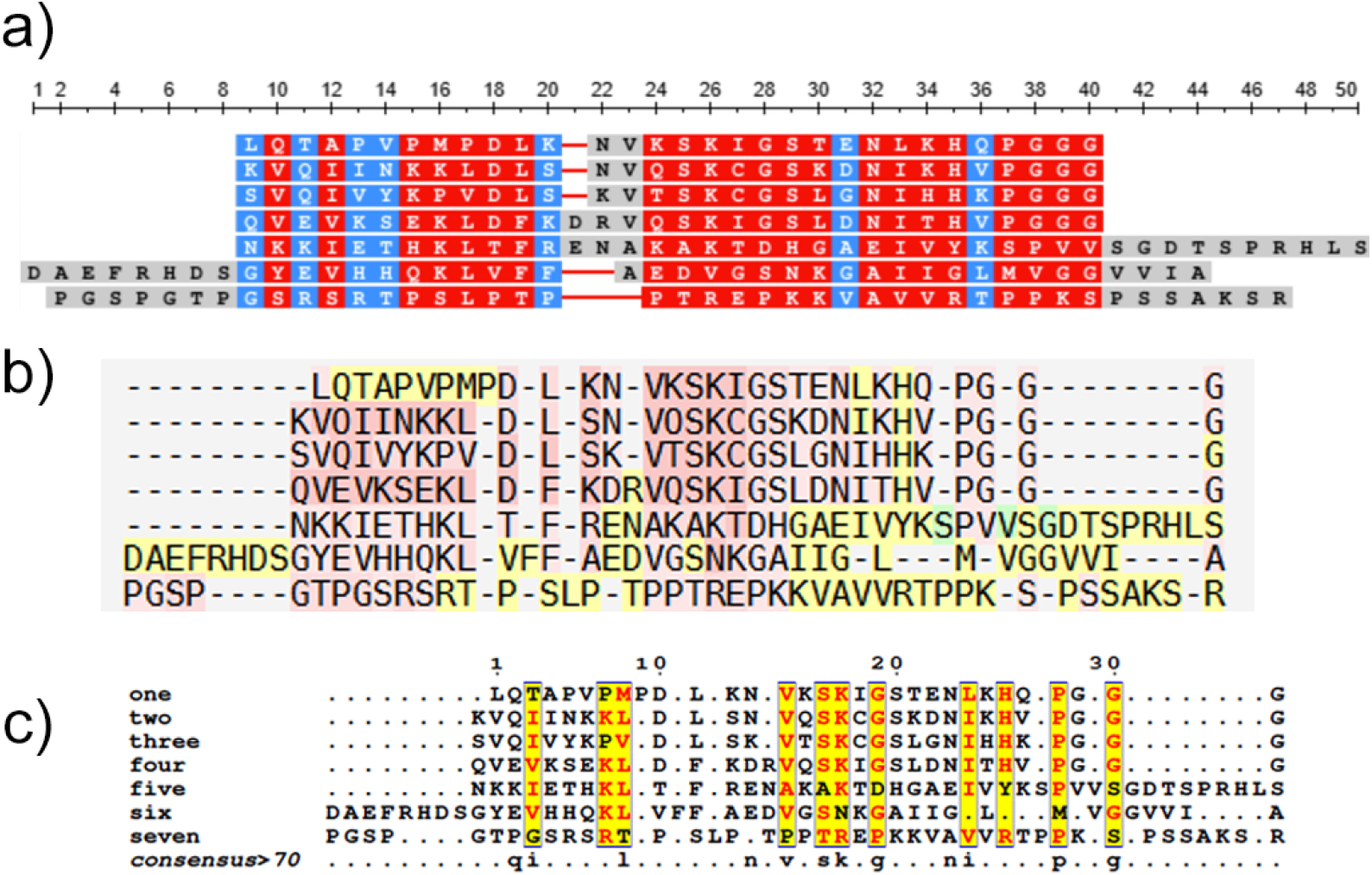
High sequence homology between Aβ1-42 and Tau R1-4, R’ and P2 domains. (a) Cobalt algorithm, red = high conservation, blue = low conservation. (b) M-Coffee algorithm, red = high alignment, yellow = average alignment. (c) Espript algorithm, highlighted = high alignment.

### Aβ binding to tubulin monomer

Initially, we examined the interaction between Aβ1-40 and Aβ1-42 with monomeric tubulin proteins by fluorescence polarization (FP) as illustrated in Figure 2. Aβ concentrations were kept at 10 nM in solution to be significantly below the aggregation critical point for either species, while tubulin concentrations were varied from 250 nM to 50 µM.^30^ The calculated K_D_ values suggest relatively weak binding between Aβ and the tubulin monomer. The binding of a single tau domain to microtubules has been shown to bridge multiple tubulin monomers,^31,32^ and given the high sequence homology, it is likely that Aβ would bridge across multiple tubulins as well. Therefore, the relatively weak binding of monomeric tubulin to Aβ may not accurately reflect the interactions between Aβ and tubulin in microtubules.

**Figure 2.**
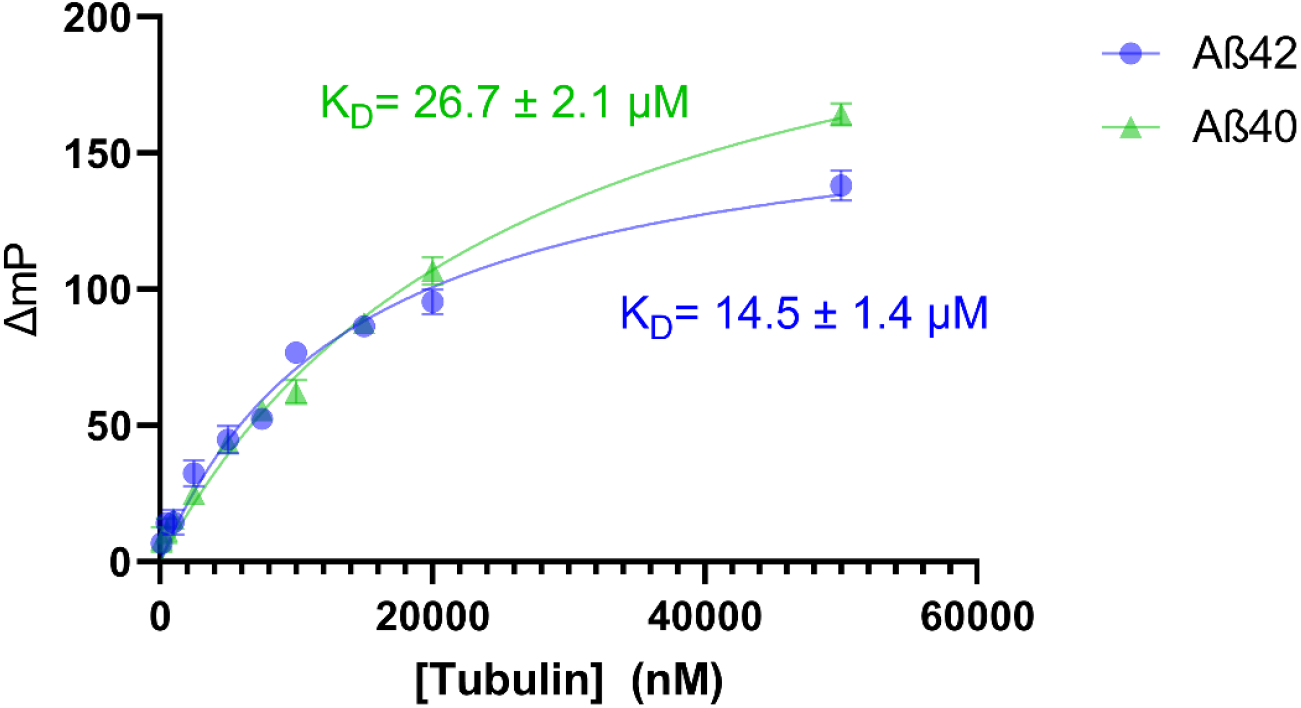
Monomeric tubulin and Aβ binding monitored by fluorescence polarization. Aβ1-40 is labeled in green, Aβ1-42 is labeled in blue.

### Amyloid beta binding to intact microtubules

To examine the binding of Aβ to microtubules, we used taxol stabilized microtubules as a model system. Taxol has been shown to bind the luminal surface of microtubules, whereas the tau binding domains interact with the outer interface between α and β tubulin heterodimers,^33-35^ therefore, taxol should not interfere with binding. Due to the relatively large size and slow diffusion expected for both Aβ and microtubules, polarization was monitored over a 24 h period to afford sufficient time for interaction. The FP results indicate K_D_ values of 2.72 ± 0.13 µM and 5.33 ± 0.83 µM for Aβ1-42 and Aβ1-40, respectively (see Fig. 3). These results confirm that Aβ does bind to microtubules with moderate affinity. Aβ1-42 shows higher binding affinity to microtubules compared to Aβ1-40 by a factor of ∼2x, which may be owed to hydrophobic interactions between the microtubules and the two additional C-terminal amino acids.

**Figure 3.**
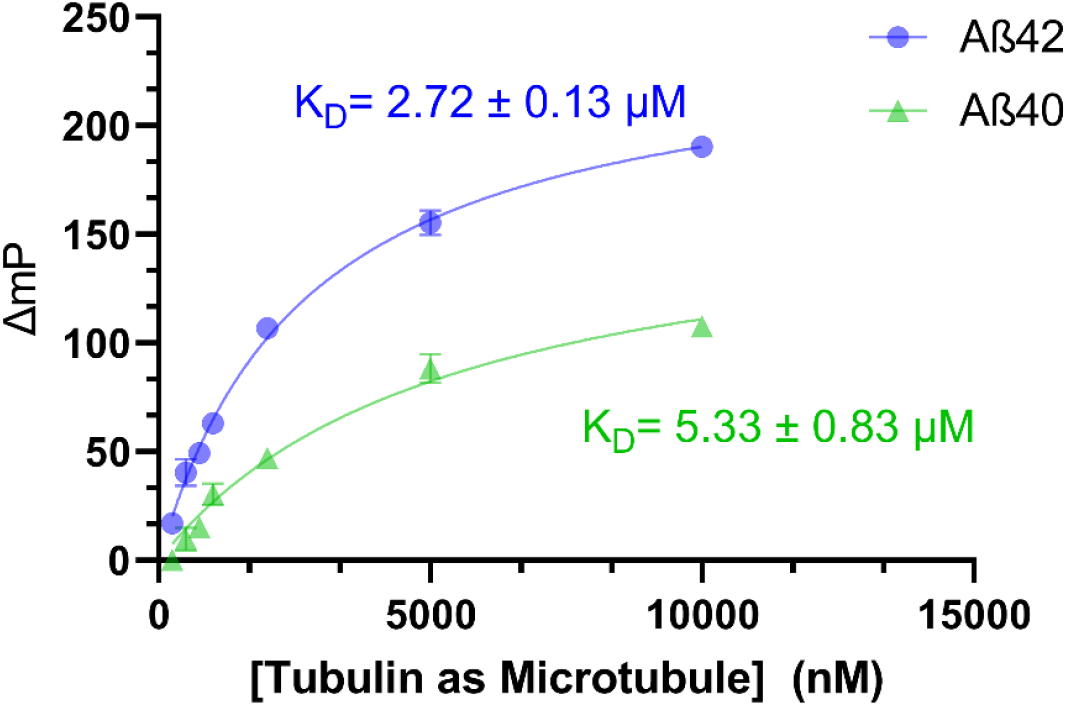
Microtubule and Aβ binding monitored by fluorescence polarization. Aβ1-40 is labeled in green, Aβ1-42 is labeled in blue.

To confirm whether Aβ and tau compete for binding sites on microtubules, we conducted competitive binding experiments monitored by FP. After allowing 24 h for Aβ-HiLyte peptides to bind to microtubules, 2 µM recombinant human tau was introduced, and FP was monitored (Fig. 4a-b). Binding to microtubules was reduced (but not eliminated) in the presence of tau for both Aβ peptides, suggesting that the affinity of tau for microtubules is comparable to that of Aβ. Furthermore, the reduction in Aβ binding implies that tau and Aβ are binding (in whole or part) to the same domains on microtubules and with similar affinities. Aβ, microtubules, and tau were also incubated simultaneously (Fig. 4c-d), and polarization was monitored as all three approached equilibrium. Under these conditions as well, tau interferes with but does not fully prevent Aβ binding. Our results are also consistent with previously reported binding affinities in the literature (see Table 1). Although there is a range of reported affinities of tau for microtubules (from mid-nanomolar to micromolar) and the previous affinity of Aβ for microtubules suggested tighter binding than our value (∼400 nM vs 2.7 μM), these variations can be attributed to differences in the experimental approaches used to obtain them. Importantly, regardless of what the true affinities are, our competition experiments test affinity in a relative fashion using a single readout in a single experiment, and the results reveal comparable affinity for tau and Aβ when binding to microtubules. Given that the binding affinities appear to be comparable, our results suggest that Aβ could inhibit tau from binding to microtubules.

**Table 1.**
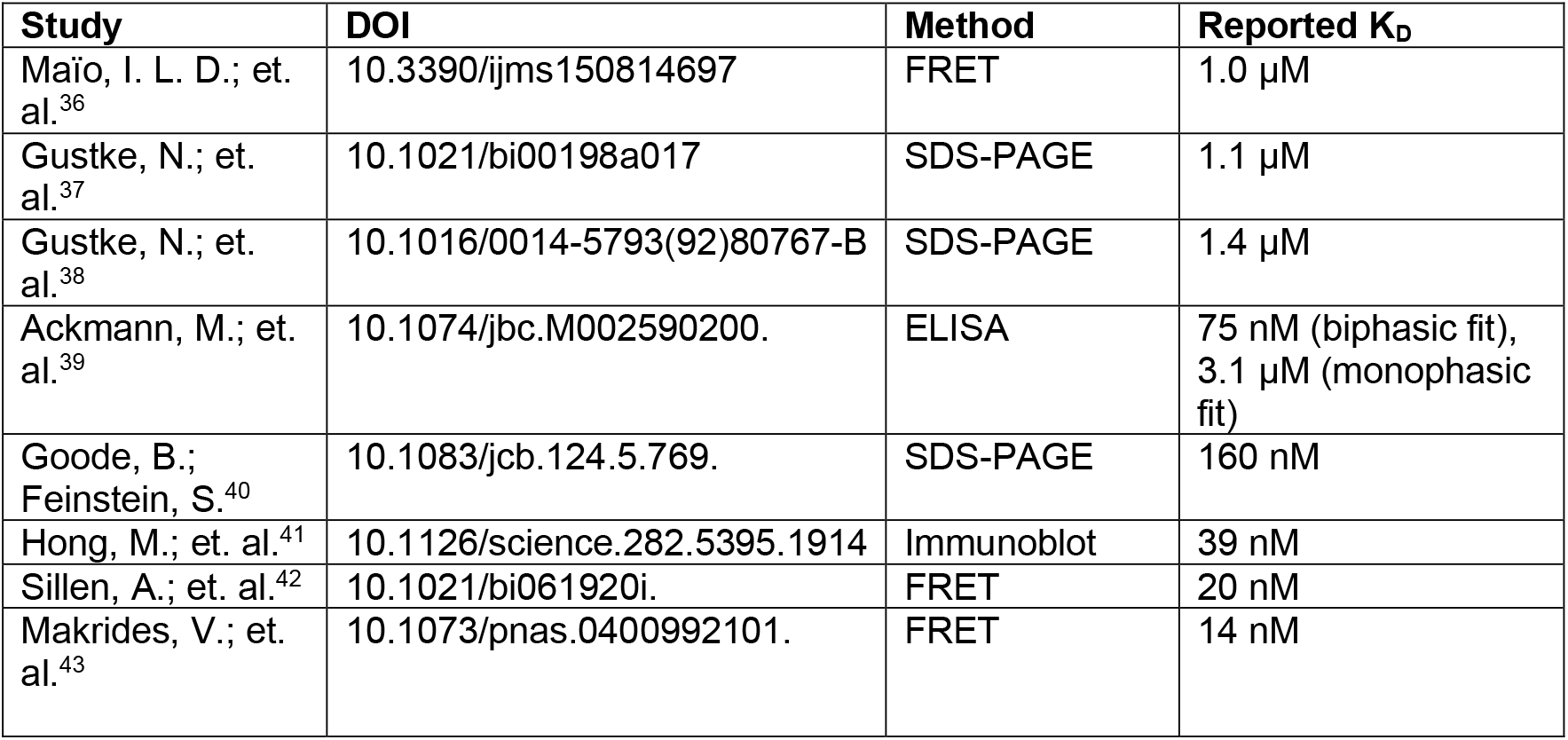
Binding affinities reported in the literature of tau for microtubules.

**Figure 4.**
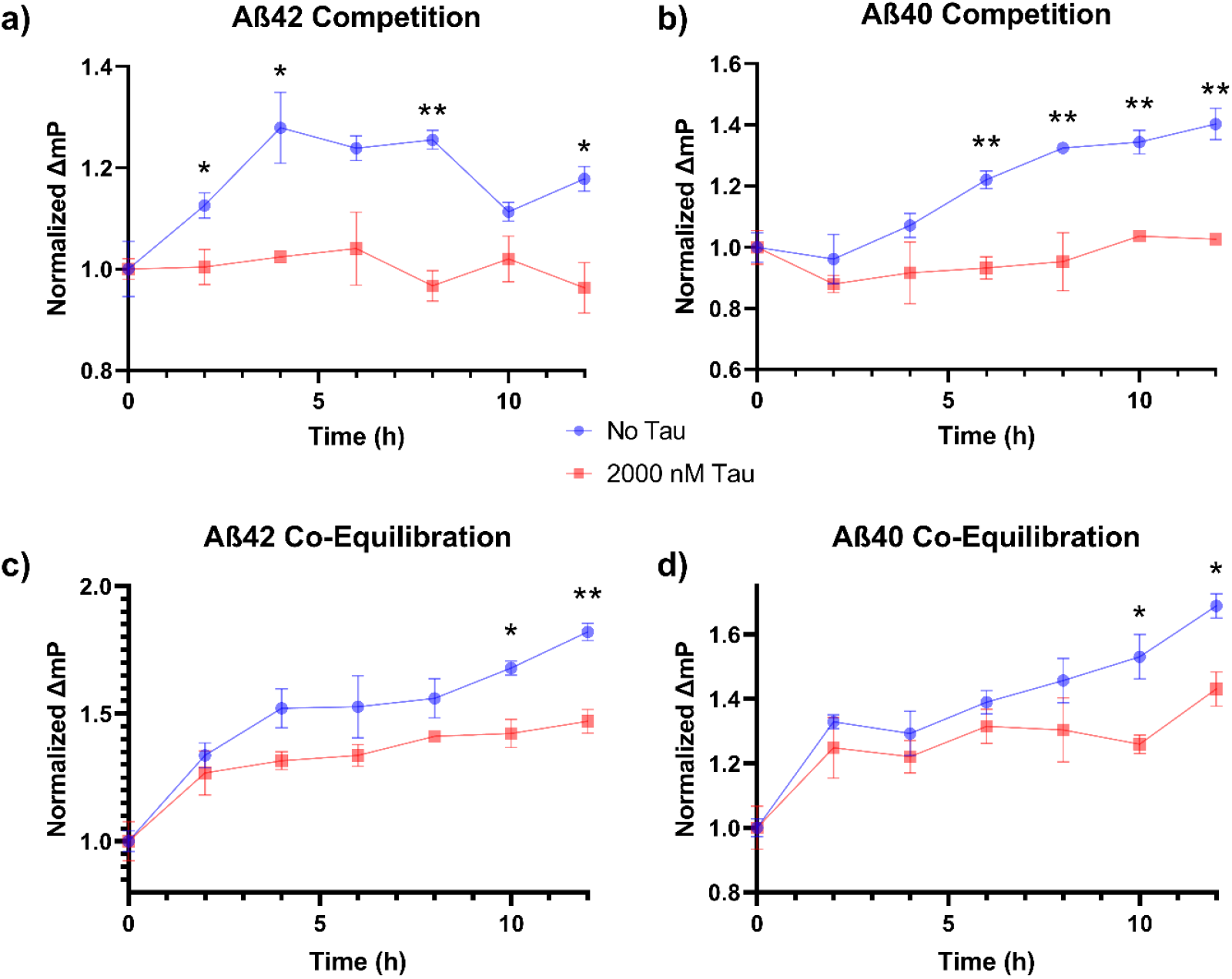
Competitive binding between tau with Aβ1-42 (a) or Aβ1-40 (b), as well as co-equilibration of microtubules, tau, and Aβ1-42 (c) or Aβ1-40 (d). Comparison between no tau added (blue) and 2000 nM tau added (red) were made using multiple unpaired t-tests assuming equal variance with the Holm-Šídák correction for multiple hypothesis testing. ^*^ = p <0.05, ^**^ = p < 0.01.

## Discussion

In order for Aβ to competitively influence tau binding to microtubules within neurons, Aβ must be produced in or relocated to the neuronal cytoplasm. Although amyloid plaques composed of Aβ are primarily found in the extracellular space between cells in the brain, and some amount of Aβ is likely generated directly in that space, several reports have concluded that the majority of Aβ originates intracellularly within neurons in the endosomal machinery.^44-46^ This finding is supported by the observation that the highest levels of the amyloid precursor protein (APP) are found within endosomes, and the BACE enzyme that catalyzes the final cleavage to create Aβ is most active at low pH, which is attained in endosomes but not in the extracellular matrix.^47^ It is also possible for extracellular Aβ to be reabsorbed into neurons through endocytosis.^48^ Therefore, there are multiple pathways by which Aβ and microtubules could become colocalized within neurons.

Competitive binding of Aβ to microtubules should interfere with the binding of tau or other proteins that bind microtubules and therefore degrade microtubule function. A variety of previous observations are consistent with this hypothesis. In studies of primary cultured neurons, addition of Aβ led to beading of microtubules within 3.5 hours. The disrupted microtubules strongly resembled those found within dystrophic neurites in amyloid plaque regions observed in brain sections of AD patients and mouse model brains.^49^ Studies using cell cultures and primary cortical neurons revealed microtubule disassembly within hours after exposure to Aβ.^50^ Substantial evidence also indicates loss of microtubule function to be a general feature of AD. For example, decreased levels of microtubules are observed in AD; however, microtubule levels don’t track with tau aggregation (suggesting that aggregation of tau does not precede microtubule pathology).^51^ Microtubules also decrease with age in cognitively normal individuals, which is consistent with the recognized age-dependence of AD (i.e. the general decrease in microtubules that occurs with age is exacerbated to pathological levels in AD).^51^ In AD model systems, microtubule shortening precedes neurodegeneration.^52^ In sum, a wealth of evidence supports the possibility that Aβ binding directly impairs the functionality of microtubules, and that this event may be an important factor in the progression of AD.

Displacement of tau from microtubules by Aβ should also modulate the behavior of tau itself. Again, many previous reports are consistent with this idea. For example, high Aβ levels lead to the delocalization of tau within neurons and increase distribution into the somatodendritic compartment.^53^ Importantly tau is an intrinsically disordered protein, particularly in the absence of binding partners.^54^ Therefore, displacement of tau from microtubules will increase the degree to which tau is disordered. This absence of structure favors aberrant hyperphosphorylation,^55^ isomerization,^56^ and aggregation^54^ of tau. These modifications in turn lower the affinity of tau for microtubules and effectively and modulate its functionality in an irreversible fashion. For example, hyperphosphorylation of tau inhibits microtubule assembly and can disrupt existing microtubules.^57^ Aβ injections were also observed to increase the rate of tau aggregation.^58^ Therefore, by preventing tau from binding to microtubules, Aβ alters the behavior of tau in ways that are pathological and include consequences that are independent of microtubule function.

Collectively, these observations suggest that the amyloid cascade hypothesis requires revision (see Scheme 1). Pathology is initiated by high levels of Aβ, which can be attributed to either increased production (e.g. many familial AD variants) or reduced clearance (e.g. due to lower autophagy inherent with aging).^59,60^ At high levels, Aβ competitively displaces some fraction of tau from microtubules. This displacement of tau is the primary toxic element of Aβ behavior. Aggregation into fibrils or oligomers is a secondary effect of the increased abundance and not a primary cause of toxicity. Displacement of tau from microtubules is also the primary source of tau toxicity. It facilitates delocalization into the somatodendritic compartment, phosphorylation, microtubule failure, and ultimately aggregation of tau and neuronal death. For tau, aggregation is an important secondary cause of toxicity that occurs in the later stages of neuronal disintegration. Importantly, displacement of tau by Aβ effectively mimics the outcome of genetic mutations to tau that interfere with microtubule binding in tauopathies, where the displacement from microtubules is recognized as the cause of the disease.^61,62^

**Scheme 1.**
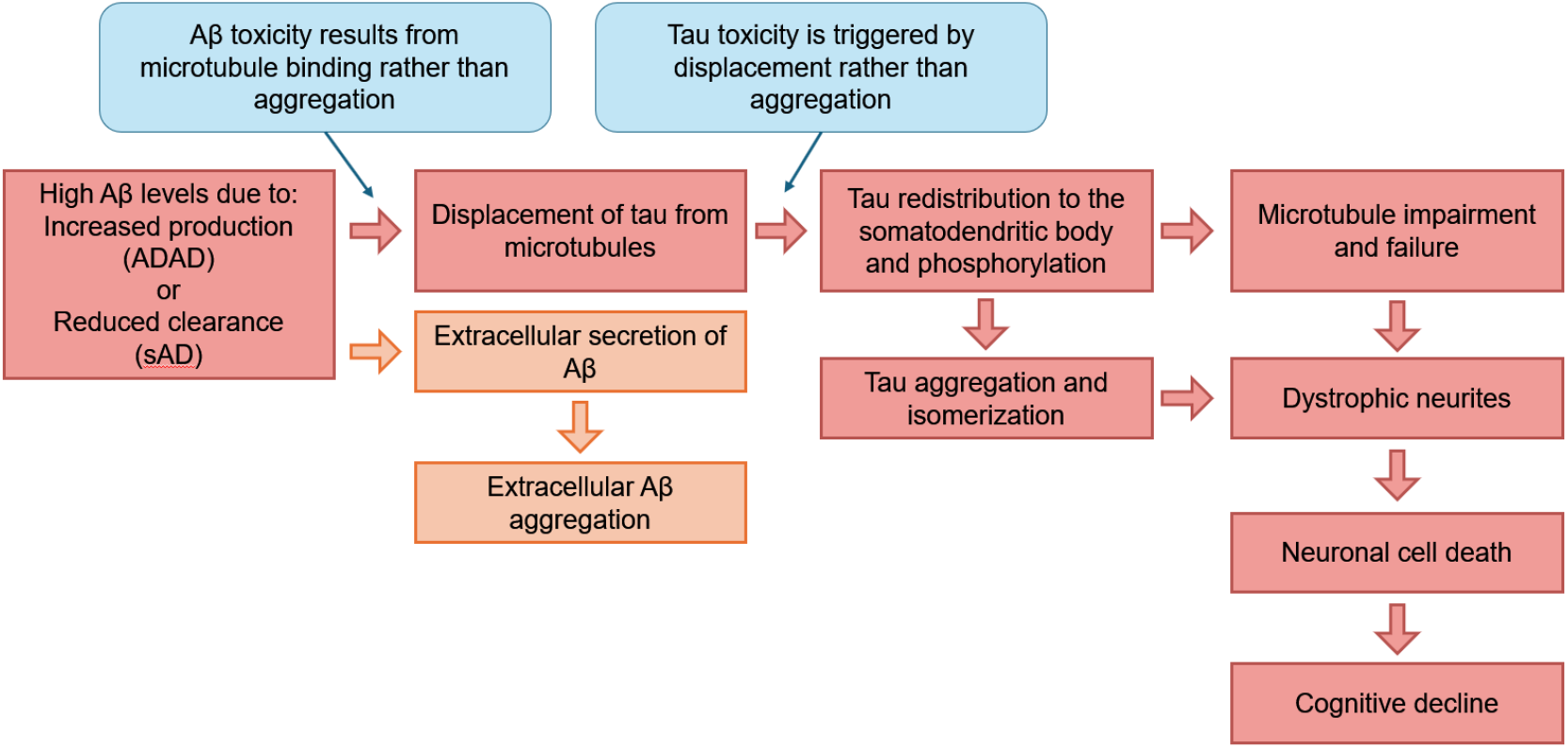
The microtubule nexus hypothesis of Alzheimer’s disease. ADAD, autosomal dominant AD; sAD, sporadic AD

The microtubule nexus hypothesis addresses many critical shortcomings that have been noted by critics of previous theories. First, Aβ and tau are both central players that directly interact with each other to induce disease pathology, thus avoiding the drawbacks of more Aβ- or tau-centric disease models. Second, amyloid plaques or oligomers of Aβ do not play a primary role, which allows for accommodation of the disconnect between the prevalence of plaques within the brain and cognitive status. To clarify, pathological displacement of tau by Aβ occurs upstream of Aβ aggregation and could therefore become problematic even in the absence of amyloid deposits. Alternatively, toxicity could be effectively averted in some neurons by efficient excretion of Aβ, leading to extracellular plaques that would in reality be indicators of protective action. On the other hand, if Aβ production simply exceeded degradation or excretion, then the presence of amyloid plaques would correlate with loss of neuronal function. In other words, the combination of these three scenarios would suggest that the simple presence or absence of Aβ plaque alone should be a very poor predictor of cognitive status (and it is). Furthermore, all of these situations likely exist in AD, and recognizing that aggregation is not the cause of toxicity allows them to be reconciled.

Third, tau is ultimately responsible for toxicity, which explains why tau markers of pathology (in particular phosphorylation and isomerization) track closely with cognitive status.^63^ The key distinction here is the recognition that tau does not initiate pathology on its own but becomes problematic after displacement by Aβ. Other factors are also easily rationalized by the microtubule nexus. For example, APOE expression is the largest genetic risk factor in sporadic AD. APOE is thought to influence Aβ trafficking, which could play an important role by increasing neuronal uptake of extracellular Aβ (thus defeating or competing with beneficial secretion).^64^ Other work has shown a protective role for lithium in AD.^65,66^ Interestingly, lithium is well-known for stabilizing microtubules,^67^ which might help counter the negative effects induced by disruption of microtubule function by Aβ binding (although it might not prevent other aspects of tau toxicity). The results and discussion presented herein do not in any way prove the microtubule nexus hypothesis is correct, but the idea certainly appears to be sufficiently compelling to merit further exploration.

## Acknowledgements

The authors gratefully acknowledge the NIH (NIA R01AG066626) for funding. We thank Yu-Hsuan Chen and Linlin Zhao for their guidance with and use of the fluorescence polarimeter.

